# Puromycin selection of cells with a high expression of the cytochrome P450 CYP3A4 gene activity from a patient with drug-induced liver injury (DILI) and their lifespan prolongation using a combination of CDK4^R24C^, cyclin D1 and TERT

**DOI:** 10.1101/2020.04.25.061275

**Authors:** Shoko Miyata, Noriaki Saku, Palaksha Kanive Javaregowda, Kenta Ite, Masashi Toyoda, Toru Kimura, Hiroshi Nishina, Atsuko Nakazawa, Mureo Kasahara, Hidenori Nonaka, Tohru Kiyono, Akihiro Umezawa

**Affiliations:** Center for Regenerative Medicine, National Center for Child Health and Development Research Institute, Tokyo, 157-8535, Japan; Department of Developmental and Regenerative Biology, Medical Research Institute, Tokyo Medical and Dental University, Tokyo, 113-8510, Japan; Department of Molecular Pathology, Tokyo Medical University, Tokyo,160-8402, Japan; Research team for Geriatric Medicine (Vascular Medicine), Tokyo Metropolitan Institute of Gerontology, Tokyo, 173-0015, Japan; Department of BioSciences, Kitasato University School of Science, Kanagawa, 252-0373, Japan; Organ Transplantation Center, National Center for Child Health and Development, Tokyo, 157-8535, Japan; Project for Prevention of HPV-related Cancer, Exploratory Oncology Research & Clinical Trial Center, National Cancer Center, Chiba 277-8577, Japan

## Abstract

Many drugs have the potential to induce the expression of drug-metabolizing enzymes, particularly cytochrome P450 3A4 (CYP3A4), in hepatocytes. Hepatocytes can accurately evaluate drug-mediated CYP3A4 induction as the gold standard for in vitro hepatic toxicology test, but their lot variation is an issue to be solved. Only a limited number of immortalized hepatocyte cells have been reported. In this study, we generated an immortalized cell expressing CYP3A4 from a patient with drug-induced liver injury (DILI). To generate DILI-derived cells with a high expression of CYP3A4, we employed a three-step approach: 1. Differentiation of DILI-induced pluripotent stem cells (DILI-iPSCs); 2. Immortalization of the differentiated cells; 3. Selection of the cells with puromycin. We hypothesize that cells with a high expression of cytochrome P450 genes can survive even after exposure to cytotoxic antibiotics because of high drug-metabolism activity. Puromycin, one of the cytotoxic antibiotics, was used in this study because of its rapid cytocidal effect at a low concentration. Phenotypic studies *in vitro* revealed that the puromycin-selected cells (HepaSM or SI cells) constitutively expressed the CYP3A4 gene at an extremely high level, and continued to proliferate at least up to 34 population doublings for more than 250 days. The expression profiles were independent of population doublings. Drug-mediated induction test revealed that the cells significantly increased CYP3A4 after exposure to rifampicin, suggesting that the immortalized cells would serve as another useful source for in vitro examination of drug metabolism and CYP3A4 induction.

## INTRODUCTION

Toxicity of low-molecular drugs has been examined in animals such as rats and mice for preclinical safety tests (The Food and Drug Administration (FDA), 2012). Primary human cells are a useful tool as an *in vitro* model for toxicity and become a golden standard in pharmaceutical *in vitro* studies for clinical prediction. However, using human cells in drug screening has its drawbacks, such as limited supply of the same lot and large variations between lots due to genetic and environmental backgrounds. To solve the lot variation issue, HepG2 is commonly used to examine hepatotoxicity because of its clonal nature. HepaRG, another hepatocyte-like clone, was established from hepatoblastoma, and has an advantage due to high induction of cytochrome p450 genes (Anthérieu et al., 2010). Technology of induced pluripotent stem cells (iPSCs) enabled us to create hepatocytes from patients with drug-induced liver injury (DILI).

Human iPSCs impact numerous medical fields including clinical therapy development, drug discovery, research on inherited diseases and studies on reprogramming of differentiated cells (De Assuncao et al., 2015; Hankowski et al., 2011; Santostefano et al., 2015; Takahashi and Yamanaka, 2006). Human iPSCs also prove valuable for toxicology testing (Anson et al., 2011; Kim et al., 2018). For example, iPSC-derived hepatocytes have been shown to serve as an *in vitro* tool for understanding drug metabolism and toxicology (Takayama et al., 2012). iPSC-derived hepatocytes or hepatocyte-like cells can be obtained from the same origin repeatedly due to immortality of iPSCs (Holmgren et al., 2014; Lu et al., 2015; Sirenko et al., 2014). Although it is expected that hepatocytes differentiated from iPSCs can be utilized in drug toxicity testing, the actual applicability of iPSC-derived hepatocytes in this context has not been well examined so far. In this study, we generated iPSCs from a pediatric patient with DILI (DILI-iPSCs), and matured DILI-iPSCs into hepatocytes (DILI-hepatocytes) or other epithelial cells that would be applicable for drug toxicity testing for a useful source for in vitro toxicology test.

Numerous human cells have successfully been immortalized. Human foreskin fibroblasts are immortalized with telomerase reverse transcriptase (TERT) alone (Bodnar et al., 1998). Human marrow stromal cells are immortalized by infecting them with TERT and the human Papillomavirus type 16-derived E6 and E7 genes because the increase in telomerase activity as a result of introduction of TERT alone is not insufficient to prolong life span of marrow stromal cells (Mori et al., 2005; Okamoto et al., 2002). Human fetal cells and myoblasts are also immortalized (Terai et al., 2005). In addition to mesenchymal stromal cells and fibroblasts, epithelial cells such as mammary cells, pancreatic ductal cells, amniotic epithelium, keratinocytes, and hepatocytes have also been immortalized with both p16/Rb inactivation by E7 and telomerase activation by E6 (Kiyono et al., 1998; Tsuruga et al., 2008). Likewise, we immortalized the puromycin-selected EpCAM-positive cells with the lentiviruses carrying the genes for TERT, CDK4R24C, and cyclin D1. The puromycin-selected immortalized cells would serve as another useful source for in vitro hepatotoxicity tests of low-molecular drugs tests.

## RESULTS

### Generation of immortalized cells from iPSCs

As starting cells for hepatic differentiation, we used iPSC-K cells that are generated from a patient with DILI (Figure 1A, B, Table 1). The differentiated cells exhibited a polygonal and/or cuboidal shape with tight cell-cell contact (Figure 1C) We employed the three steps to obtain cells with an extended life span (Figure 1A). We first differentiated iPSC-K cells into endodermal cells. We selected EpCAM-positive differentiated cells with the magnetic-activated cell sorting. We then infected the lentivirus carrying combinations of the TERT, CDK4^R24C^, and cyclin D1 genes to extend a lifespan. The infected cells showed a mixed cell population in morphology (Figure 1D) and became positive for CYP3A4 in part (Figure 1E). The EpCAM-positive clone produced E-cadherin and cyclin D1 (Figure 1F).

**Table 1.** A list of immortalized cells from DILI patients.

| Table 1. A list of immortalized cells from DILI patients |  |  |  |  |  |
| --- | --- | --- | --- | --- | --- |
| Cell Name | Gene-1 | Gene-2 | Gene-3 | Gene-4 | Parental Cell |
| iHep2-D9-81 | CSII-CMV-hCDK4R24C | CSII-CMV-cyclin D1 | CSII-CMV-TERT |  | iPSC-I or Cell-I |
| iHep2-D9-85 | CSII-ALB-tetOff-ADV | CSII-TRE-Tight-cyclin D1 | CSII-TRE-Tight-hCDK4R24C | CSII-CMV-TERT | iPSC-I or Cell-I |
| iHep3-D9-89 | CSII-CMV-hCDK4R24C | CSII-CMV-cyclin D1 | CSII-CMV-TERT |  | iPSC-O or Cell-O |
| iHep3-E1-29 | CSII-ALB-tetOff-ADV | CSII-TRE-Tight-cyclin D1 | CSII-TRE-Tight-hCDK4R24C | CSII-CMV-TERT | iPSC-O or Cell-O |
| iHep2-E2-64 | CSII-CMV-hCDK4R24C | CSII-CMV-cyclin D1 | CSII-CMV-TERT |  | iPSC-I or Cell-I |
| iHep2-E2-66 | CSII-ALB-tetOff-ADV | CSII-TRE-Tight-hCDK4R24C | CSII-TRE-Tight-cyclin D1 | CSII-CMV-TERT | iPSC-I or Cell-I |
| iHep3-E2-68 | CSII-CMV-hCDK4R24C | CSII-CMV-cyclin D1 | CSII-CMV-TERT |  | iPSC-O or Cell-O |
| iHep3-E2-70 | CSII-ALB-tetOff-ADV | CSII-TRE-Tight-hCDK4R24C | CSII-TRE-Tight-cyclin D1 | CSII-CMV-TERT | iPSC-O or Cell-O |
| iHep2-E3-32 | CSII-CMV-hCDK4R24C | CSII-CMV-cyclin D1 | CSII-CMV-TERT |  | iPSC-I or Cell-I |
| iHep2-E3-34 | CSII-ALB-tetOff-ADV | CSII-TRE-Tight-cyclin D1 | CSII-TRE-Tight-hCDK4R24C | CSII-CMV-TERT | iPSC-I or Cell-I |
| iHep2-E3-36 | CSII-CMV-hCDK4R24C | CSII-CMV-cyclin D1 | CSII-CMV-TERT |  | iPSC-I or Cell-I |
| iHep3-E3-38 | CSII-CMV-hCDK4R24C | CSII-CMV-cyclin D1 | CSII-CMV-TERT |  | iPSC-O or Cell-O |
| iHep3-E3-40 | CSII-ALB-tetOff-ADV | CSII-TRE-Tight-cyclin D1 | CSII-TRE-Tight-hCDK4R24C | CSII-CMV-TERT | iPSC-O or Cell-O |
| iHep3-E3-42 | CSII-CMV-hCDK4R24C | CSII-CMV-cyclin D1 | CSII-CMV-TERT |  | iPSC-O or Cell-O |
| iHep2-E3-62 | CSII-ALB-tetOff-ADV | CSII-TRE-Tight-hCDK4R24C | CSII-TRE-Tight-cyclin D1 | CSII-CMV-TERT | iPSC-I or Cell-I |
| iHep3-E3-64 | CSII-ALB-tetOff-ADV | CSII-TRE-Tight-hCDK4R24C | CSII-TRE-Tight-cyclin D1 | CSII-CMV-TERT | iPSC-O or Cell-O |
| iHep4-E4-72 | CSII-CMV-hCDK4R24C | CSII-CMV-cyclin D1 | CSII-CMV-TERT |  | iPSC-K or Cell-K |
| iHep4-E4-74 | CSII-ALB-tetOff-ADV | CSII-TRE-Tight-cyclin D1 | CSII-TRE-Tight-hCDK4R24C | CSII-CMV-TERT | iPSC-K or Cell-K |
| iHep4-E4-80 | CSII-CMV-TERT | CSII-CMV-hCDK4R24C | CSII-CMV-cyclin D1 |  | iPSC-K or Cell-K |
| iHep4-E4-82 | CSII-CMV-TERT | CSII-CMV-hCDK4R24C | CSII-CMV-cyclin D1 |  | iPSC-K or Cell-K |
| iHep4-E4-84 | CSII-ALB-tetOff-ADV | CSII-TRE-Tight-cyclin D1 | CSII-TRE-Tight-hCDK4R24C | CSII-CMV-TERT | iPSC-K or Cell-K |
| iHep4-E5-50 | CSII-CMV-TetOff-ADV | CSII-TRE-Tight-cyclin D1 | CSII-TRE-Tight-hCDK4R24C | CSII-CMV-TERT | iPSC-K or Cell-K |
| iHep2-E8-45 | CSII-ALB-tetOff-ADV | CSII-TRE-Tight-hCDK4R24C | CSII-TRE-Tight-cyclin D1 | CSII-CMV-TERT | iPSC-I or Cell-I |
| iHep2-E8-46 | CSII-ALB-tetOff-ADV | CSII-TRE-Tight-hCDK4R24C | CSII-TRE-Tight-cyclin D1 | CSII-CMV-TERT | iPSC-I or Cell-I |
| iHep4-E8-49 | CSII-CMV-TetOff-ADV | CSII-TRE-Tight-cyclin D1 | CSII-TRE-Tight-hCDK4R24C | CSII-CMV-TERT | iPSC-K or Cell-K |
| iHep4-E8-50 | CSII-CMV-TetOff-ADV | CSII-TRE-Tight-cyclin D1 | CSII-TRE-Tight-hCDK4R24C | CSII-CMV-TERT | iPSC-K or Cell-K |

**Figure 1.**
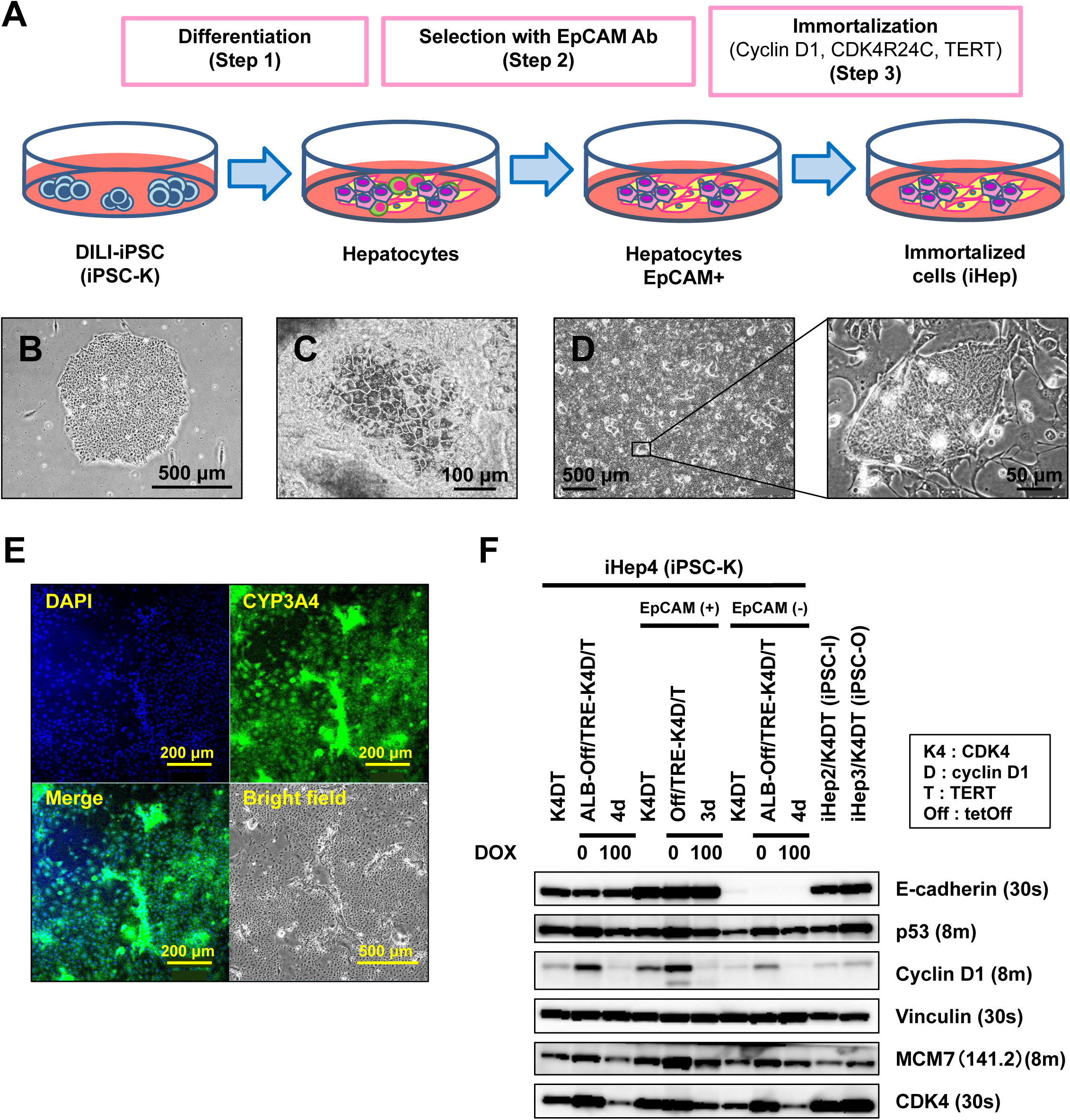
Immortalization of hepatocytes from iPSCs. A. Scheme for generation of immortalized hepatocytes from iPSCs.Hepatocyte cells are induced from iPSCs (step 1). To immortalize hepatocyte cells, EpCAM-positive cells are sorted from iPSC-derived hepatocytes (step 2) are infected with the lentiviral vectors carrying TERT, CDK4^R24C^, and cyclin D1 (step 3). B. Phase-contrast photomicrograph of iPSC-K. iPSC-K cells on MEFs formed a flat and well-defined colony. C. Phase-contrast photomicrograph of Hepatocyte-like cells from iPSC-K. D. Phase-contrast micrograph of immortalized iHep4 cells. E. Immunocytochemistry of iHep4 cells using the antibodies to CYP3A4. F. Western blot analysis of iHep4 cells using the antibodies to E-cadherin, P53, cyclin D1, vinculin, MCM7, and CK7.

### Puromycin selection of the differentiated cells from liver injury-derived iPSC-K

Puromycin is an antibiotic that is toxic to eukaryotic cells and can be metabolized by cytochrome P450. To isolate cells with a high expression of the CYP3A4 gene, from the EpCAM-positive cells, we exposed the cells to puromycin for 3 days (Figure 2A). Through this selection method, we successfully obtained a population of cells with a parenchymal or epithelial cell morphology (Figure 2B, C). The selected cells exhibited endodermal cell morphologies, i.e., normal hepatocyte- or intestinal cell-morphology: Clear distinct one or two round nuclei with prominent nucleoli and dark cytoplasm, and confluent compact monolayer colonies.

**Figure 2.**
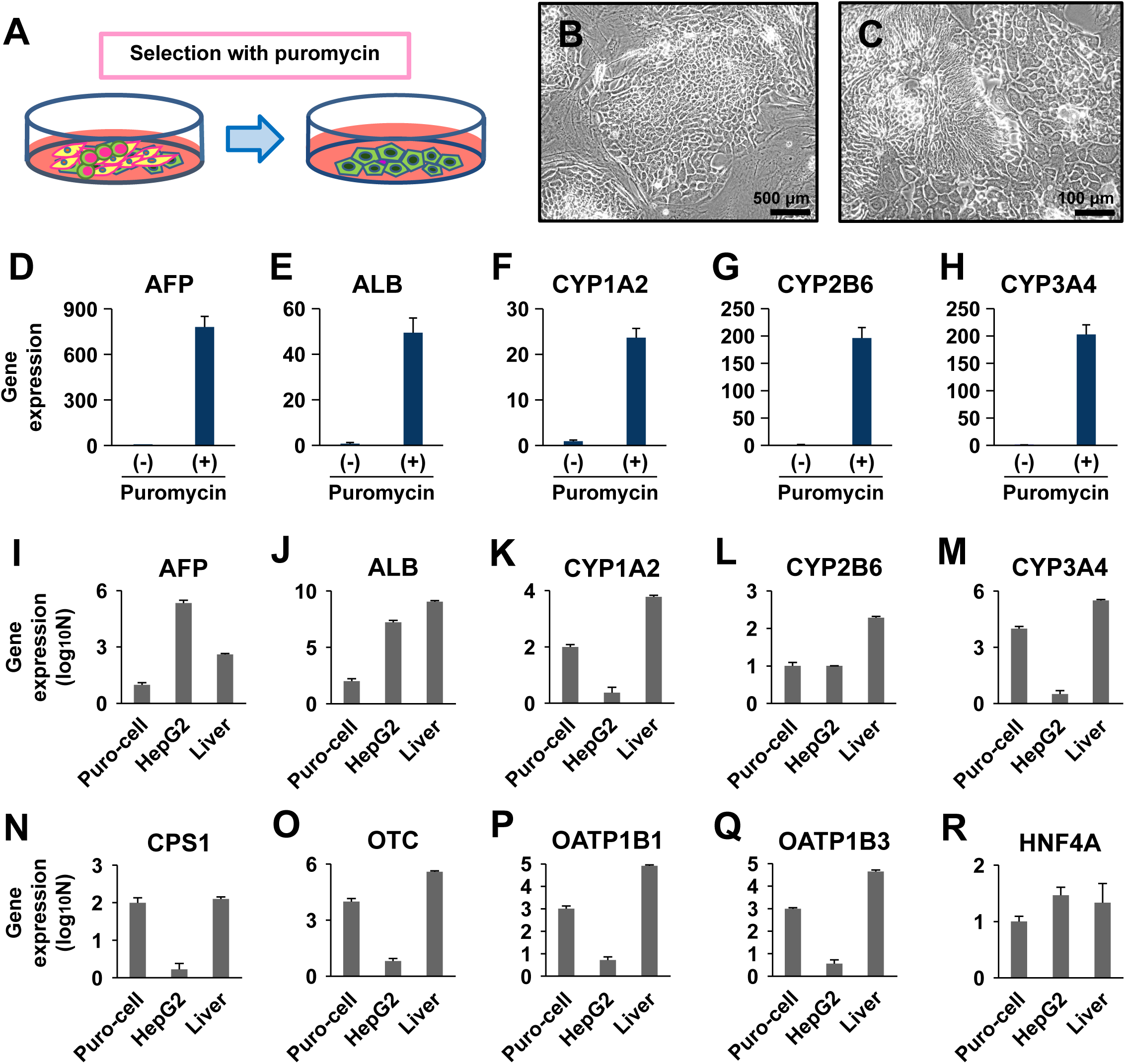
Selection of puromycin-resistant cells. A. Scheme for selection of puromycin-resistant cells (green) from a mixed population of immortalized cells. B. Phase contrast photomicrograph of the puromycin-selected cells. Low-power view. C. Phase contrast photomicrograph of the puromycin-selected cells. High-power view. D-H. Quantitative RT-PCR analysis of the genes for α-FETOPROTEIN (D: AFP), ALBUMIN (E: ALB), CYP1A2 (F), CYP2B6 (G), and CYP3A4 (H) after the puromycin treatment. The immortalized differentiated cells (Figure 1A) were exposed to puromycin at a final concentration of 3 μg/ml for 3 days. RNAs were isolated from the cells at one week after the puromycin treatment. (-): no puromycin treatment, (+): puromycin treatment. mRNA levels were normalized using UBIQUITIN as a housekeeping gene. The amounts of each gene at the cells without puromycin treatment were regarded as equal to 1. I-R. Quantitative RT-PCR analysis of the puromycin-selected immortalized cells after propagation (Puro-cell), HepG2 cells, and hepatocytes (Liver). I: α-FETOPROTEIN (AFP), J: ALBUMIN (ALB), K:CYP1A2, L:CYP2B6, M:CYP3A4, N: CARBAMOYL PHOSPHATE SYNTHETASE 1 (CPS1), O: ORNITHINE TRANSCARBAMYLASE (OTC), P:OATP1B1, Q:OATP1B3, R:HNF4A. mRNA levels were normalized using UBIQUITIN as a housekeeping gene. Relative expression levels of each gene were shown.

### Changes in gene expression by puromycin selection

In order to verify whether drug selection with puromycin influences gene expression, we performed qRT-PCR analysis. The puromycin-selected cells exhibited higher expression of the genes for AFP, ALB, CYP1A2, CYP2B6, and CYP3A4 (Figure 2D-H). Ratio of the cells with hepatocyte- or intestinal epithelium-like morphology was increased in morphology after the puromycin treatment. We then compared expressions of each gene in the puromycin-selected cells with HepG2 cells and hepatocytes (Figure 2I-R). The puromycin-selected cells expressed a lower level of the AFP gene than HepG2 cells, and a lower level of the ALB genes than the hepatocyte sample; the puromycin-selected cells expressed a comparable level of the genes for CYP1A2, CYP2B6, CYP3A4, CPS1, OTC, OATP1B1, OATP1B3, and HNF4A, compared with the hepatocyte sample.

### Characterization of the puromycin-selected cells

To investigate characterization of the puromycin-selected cells further, we performed karyotype analysis, cytochrome P450 induction test, and long-term growth analysis. The karyotypic analysis showed that the puromycin-selected cells had intact chromosomes except for chromosome 8 trisomy in the cell stock (Figure 3A). The puromycin-selected cells was immunocytochemically positive for AFP, ALB, and CYP3A4 (Figure 3B, C). The puromycin-selected cells in iPGell displayed an eosinophilic cytoplasm and basophilic stippling, and round nuclei with dispersed chromatin in a form of gel as live cells (Figure 3D). Ultrastructural analysis revealed desmosome at the cell junction and microvilli on the cell surface (Figure 3E, F). The puromycin-selected cells were positive for CK8/18 (AE1/3) (Figure 3G, H), E-cadherin (Figure 3I, J), PCNA (Figure 3K, L) and Ki67 (Figure 3M, N). We then evaluated the puromycin-selected cells for cytochrome P450 induction. We investigated expression levels of major three cytochrome P450 enzymes, CYP1A2, CYP2B6 and CYP3A4 (Figure 3O-Q). We exposed the puromycin-selected cells to omeprazole for 24 h, phenobarbital for 48 h, and rifampicin for 48 h. Expression of the CYP1A2 gene was up-regulated at 3.7 fold with exposure to omeprazole (Figure 3O); expression of the CYP2B6 gene was up-regulated at 1.9 fold with exposure to phenobarbital (Figure 3P); expression of the CYP3A4 gene was up-regulated at 2.0 fold with exposure to rifampicin (Figure 3Q). Long-term cultivation analysis revealed that the puromycin-selected cells continued to proliferate up to at least 34 population doublings for more than 270 days (Figure 3R).

**Figure 3.**
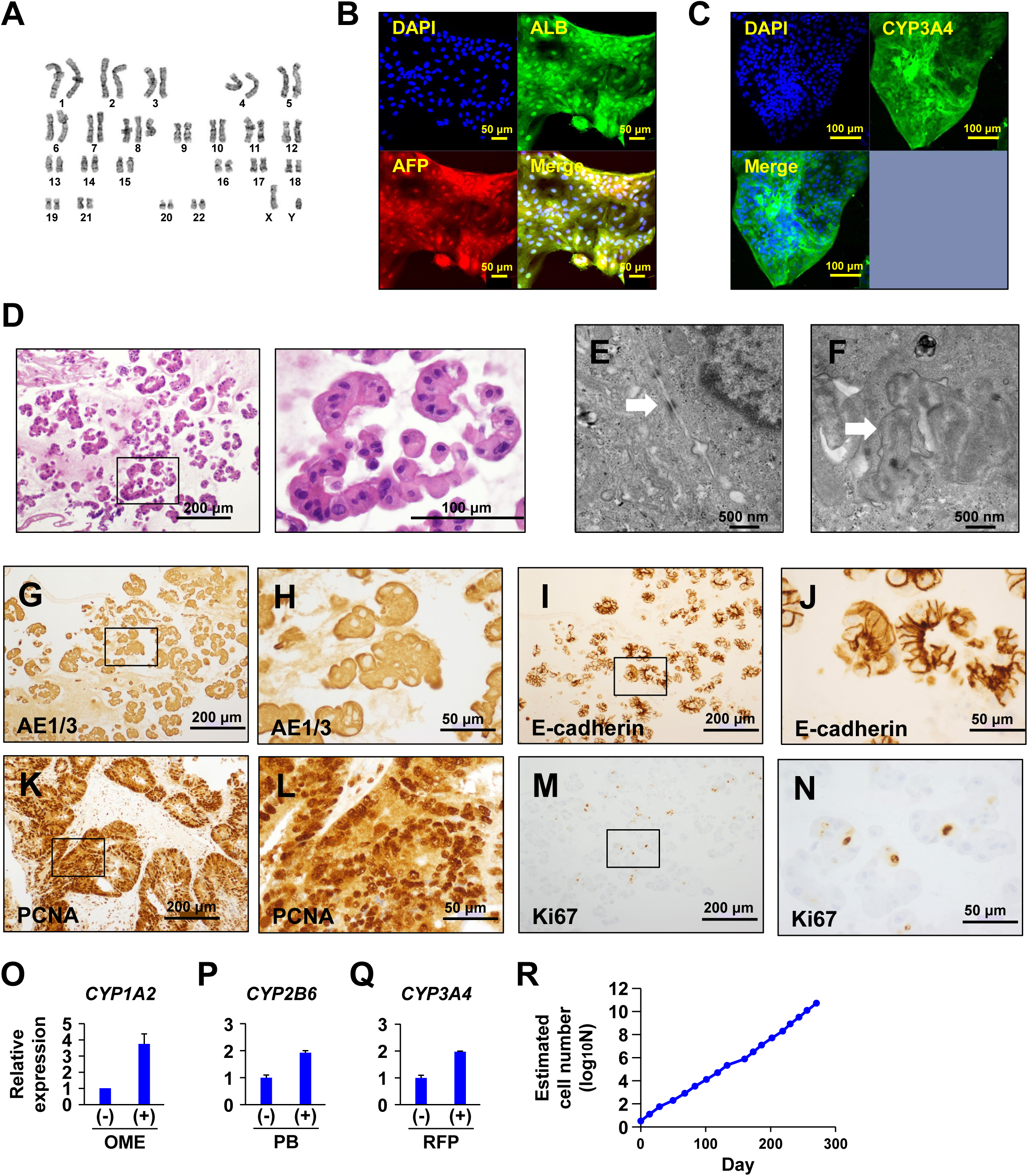
Characterization of the puromycin-selected cells. A. Karyotypic analysis of the puromycin-selected cells. B. Immunocytochemistry of the puromycin-selected cells with the antibodies to ALB (green) and AFP (red). C. Immunocytochemistry of the puromycin-selected cells with the antibodies to CYP3A4 (green). D. Histological analysis of the puromycin-selected cells embedded in iPGell. The puromycin-selected cells exhibited hepatocyte-like morphology. HE stain. E&F. Ultrastructural analysis. E: Desmosome (arrow), F: Microvilli (arrow). G-N. Immunocytochemistry of the puromycin-selected cells with the antibodies to AE1/AE3 (G, H), E-cadherin (I, J), PCNA (K, L), and Ki67 (M, N). Panel H, J, L, and N are high-power views of Panel G, I, K, and M, respectively. O. Quantitative RT-PCR analysis of the CYP1A2 gene on the puromycin-selected immortalized cells with exposure to 50 μM omeprazole (OME) for 24 h. mRNA levels were normalized using ubiquitin expression as a housekeeping gene. P. Quantitative RT-PCR analysis of the CYP2B6 gene on the puromycin-selected immortalized cells with exposure to 500 μM phenobarbital (PB) for 48 h. Q. Quantitative RT-PCR analysis of the CYP3A4 gene on the puromycin-selected immortalized cells with exposure to 20 μM rifampicin (RFP) for 48 h. R. Growth curve of the puromycin-selected immortalized cells.

## DISCUSSION

In this study, we established the puromycin-selected cells with high expression of CYP3A4 from a patient with DILI. The cells with drug-mediated CYP3A4 induction potency as well as HepG2, HepaRG and hepatocytes may serve a model cell in systems such as multi-physiological system, organ/organoid on a chip or body on a chip for examination of drug metabolism and toxicity (Park et al., 2019). We have previously immortalized human mesenchymal stromal cells derived from bone marrow by the introduction of Simian Vacuolating Virus 40 large T antigen, Polyomavirus middle T antigen or human TERT with human Papillomavirus type 16-derived E6/E7 protein (Cui et al., 2007; Mori et al., 2005; Terai et al., 2005; Umezawa et al., 1992). The difficulty, i.e. non-proliferative nature of the iPSC-derived differentiated cells, was solved by the introduction of human Cyclin D1, mutant CDK4, and TERT. The version of CDK4 used in this study has a mutation (R24C) at the binding site for the senescence protein p16 (Sasaki et al., 2009; Yugawa et al., 2007; Zushi et al., 2011). This method of cell immortalization was initially reported in humans and other animals (Donai et al., 2014; Kuroda et al., 2015; Sasaki et al., 2009). The DILI-iPSC-derived cells immortalized with the introduction of the three genes, i.e. human Cyclin D1, mutant CDK4, and telomerase, is another successful example for establishment of a useful cell with stable proliferation and genetic stability. There have been no reports of a sufficient number of DILI-derived cells. One reason is that hepatocytes are injured in patients with DILI and could not thus be obtained. Another is that hepatocytes undergo a limited number of divisions in culture and then enter a nondividing state referred to as “senescence” (Allsopp et al., 1992; Bodnar et al., 1998; Wright and Shay, 1992).

From the viewpoint of the extremely high CYP3A4 level, the puromycin-selected immortalized cells are considered one of the model cells for in vitro drug toxicology test. Because CYP3A4 contributes to the first-pass metabolism of many commercial drugs, it is important to investigate the CYP3A4-mediated hepatic metabolism in order to estimate hepatotoxicity. It is known that the CYP3A4 expression can be induced by various drugs, such as dexamethasone, PB, RFP, and 1α,25-dihydroxyvitamin D3. The induction of CYP3A4 expression by such drugs might affect the pharmacokinetics of concomitant drugs administered orally. This cell model with reproducible induction of CYP3A4 would, therefore, be useful for an early stage of drug development. Another advantage of this cell model is unchanged during in vitro propagation for banking and thus available for cytochrome p450 induction tests and cell toxicity examination, suggesting the robust system of toxicology test and the better cost-performance balance.

## MATERIALS AND METHODS

### Ethical statement

Human cells in this study were performed in full compliance with the Ethical Guidelines for Medical and Health Research Involving Human Subjects (Ministry of Health, Labor, and Welfare, Japan; Ministry of Education, Culture, Sports, Science and Technology, Japan). Animal experiments were performed according to protocols approved by the Institutional Animal Care and Use Committee of the National Center for Child Health and Development.

### Cells

The differentiated cells from DILI-iPSCs were immortalized by infection with the lentiviral vector plasmids CSII-CMV-Tet-Off, CSII-TRE-Tight-cyclin D1, SII-TRE-Tight-CDK4^R24C^, and CSII-TRE-Tight-TERT (Shiomi et al., 2011; Yugawa et al., 2007). In brief, the EF1α promoter in CSII-EF-RfA (a gift from Dr. H. Miyoshi, RIKEN) was replaced with a tetracycline-inducible promoter, TRE-Tight, from pTRE-Tight (Clontech, Mountain View, CA) to generate CSII-TRE-Tight-RfA. Human cyclin D1, human mutant CDK4 (CDK4^R24C^: an INK4a-resistant form of CDK4), and TERT were inserted into the entry vector via a BP reaction (Invitrogen, Carlsbad, CA). These segments were then recombined with CSII-TRE-Tight-RfA through an LR reaction (Invitrogen) to generate CSII-TRE-Tight-cyclin D1, CSII-TRE-Tight-CDK4^R24C^, and CSII-TRE-Tight-TERT. The rtTA segment from pTet-Off Advanced (Clontech) was amplified by PCR, recombined with the donor vector pDONR221 via a BP reaction (Invitrogen) to generate pENTR221-Tet-Off, and then recombined with a lentiviral vector, CSII-CMV-RfA, through an LR reaction (Invitrogen) to generate CSII-CMV-Tet-Off. Recombinant lentiviruses with vesicular stomatitis virus G glycoprotein were produced as described previously (Miyoshi, 2004). EpCAM-positive cells were isolated as described below after the immortalization from a mixed population. The immortalized cells were maintained in the modified F-medium at 37°C in a humidified atmosphere containing 95% air and 5% CO2 (Yachida et al., 2016). When the cultures reached subconfluence, the cells were harvested with Trypsin-EDTA Solution (cat#23315, IBL CO., Ltd, Gunma, Japan), and re-plated at a density of 5 × 10^5^ cells in a 100-mm dish. Medium changes were carried out twice a week thereafter.

### Isolation of EpCAM-positive cells

EpCAM-positive cells were isolated from the differentiated endodermal cells using a magnetic cell sorting kit (MACS; Miltenyi Biotec K.K. Cologne, Germany) with the CD326 (EpCAM) MicroBeads (cat# 130-061-101, Miltenyi Biotec), according to the manufacturer’s instructions. The magnetically labeled EpCAM-positive cells were eluted as a positively selected cell fraction.

### Immunoblot analysis

Whole-cell protein extracts were used for analysis, and immunoblotting was conducted as described previously (Yugawa et al., 2007). Antibodies against E-cadherin (Mouse MAb IgG2a; BD Transduction Lab; BD-610181), p53 (MAb clone DO-1;IgG2a; Calbiochem (Oncogene Science) Ab6; OP43-100UG), Cyclin D1 (MAb clone G124-326; IgG1; BDPharmingen; 554180), Vinculin (mouse IgG1; SIGMA-ALDRICH; V9264), MCM7 (Mouse MAb IgG1; Santa Cruz; sc-9966), CDK4 (Rabbit Pab; Cell Signalling; #610147), were used as probes, and horseradish peroxidase-conjugated anti-mouse or anti-rabbit immunoglobulins (Jackson Immunoresearch Laboratories, West Grove, PA) were employed as secondary antibodies. The LAS3000 charge-coupled device (CCD) imaging system (Fujifilm Co. Ltd., Tokyo, Japan) was employed for detection of proteins visualized by Lumi-light Plus Western blotting substrate (Roche, Basel, Switzerland).

### Immunocytochemical analysis

Cells were fixed with 4% paraformaldehyde in PBS for 10 min at 4°C. After washing with PBS and treatment with 0.2% Triton X in PBS for 10 min, cells were pre-incubated with blocking buffer (10% goat serum in PBS) for 30 min at room temperature, and then exposed to primary antibodies to Albumin (CEDARLANE, CLFAG2140, diluted at 1/50), α-fetoprotein (R&D Systems, MAB1368, diluted at 1/100), and CYP3A4 (SANTA CRUZ BIOTECHNOLOGY, sc-53850, diluted at 1/200) in blocking buffer overnight at 4°C. Following washing with 0.2% PBST, cells were incubated with secondary antibodies; either anti-rabbit or anti-mouse IgG conjugated with Alexa 488 or 546 (1:300) (Invitrogen) in blocking buffer for 30 min at room temperature. Then, the cells were counterstained with DAPI.

### Karyotypic analysis

Karyotypic analysis was contracted out to Nihon Gene Research Laboratories Inc. (Sendai, Japan). Metaphase spreads were prepared from cells treated with 100 ng/mL of Colcemid (Karyo Max, Gibco Co. BRL) for 6 h. The cells were fixed with methanol:glacial acetic acid (2:5) three times, and placed onto glass slides (Nihon Gene Research Laboratories Inc.). Chromosome spreads were Giemsa-banded and photographed. A minimum of 10 metaphase spreads were analyzed for each sample, and karyotyped using a chromosome imaging analyzer system (Applied Spectral Imaging, Carlsbad, CA).

### Quantitative RT-PCR

RNA was extracted from cells using the ISOGEN (NIPPON GENE). An aliquot of total RNA was reverse transcribed using an oligo (dT) primer (SuperScript TM LJ First-Strand Synthesis System, Invitrogen). For the thermal cycle reactions, the cDNA template was amplified (QuantStudio TM 12K Flex Real-Time PCR System) with gene-specific primer sets (Supplemental Table 1) using the Platinum Quantitative PCR SuperMix-UDG with ROX (11743-100, Invitrogen) under the following reaction conditions: 40 cycles of PCR (95°C for 15 s and 60°C for 1 min) after an initial denaturation (50°C for 2 min and 95°C for 2 min). Fluorescence was monitored during every PCR cycle at the annealing step. The authenticity and size of the PCR products were confirmed using a melting curve analysis (using software provided by Applied Biosystems) and gel analysis. mRNA levels were normalized using ubiquitin as a housekeeping gene. cDNAs from primary hepatocytes were purchased from TAKARA Shuzo, Ltd (CLN636531 Z6531N).

### Histological analysis

Cells were harvested with a cell scraper and collected into tubes. The cells were analyzed with an iPGell kit (GenoStaff, Tokyo, Japan).

### CYP3A4 induction test

The puromycin-selected immortalized cells were treated with 50 μM omeprazole for 24 h, with 500 μM phenobarbital (PB, Wako) for 48 h, or 20 μM rifampicin (RFP, Wako) for 48 h. Controls were treated with DMSO (final concentration 0.2%).

### Statistical analysis

Statistical analysis was performed using the unpaired two-tailed Student’s t test.

## Funding information

This research was supported by the Grant of National Center for Child Health and Development and Japan Agency for Medical Research and Development and AMED. Computation time was provided by the computer cluster HA8000/RS210 at the Center for Regenerative Medicine, National Center for Child Health and Development Research Institute.

### Acknowledgements

We would like to express our sincere thanks to K. Miyado for fruitful discussion, M. Ichinose for providing expert technical assistance, C. Ketcham for English editing and proofreading, and E. Suzuki and K. Saito for secretarial work.

## Author Contribution Statement

AU designed experiments. SM, KI, TKiy, and NS performed experiments. SM and AU analyzed data. AN, MK, and TKiy contributed reagents, materials and analysis tools. PKJ, MT, TKim, HNi, HNo, and TKiy discussed the data and manuscript. AU and SM wrote this manuscript.

