## Supplemental Table 1 for "Puromycin selection of cells with a high expression of the cytochrome P450 CYP3A4 gene activity from a patient with drug-induced liver injury (DILI) and their lifespan prolongation using a combination of CDK4^R24C^, cyclin D1 and TERT"

### Supplemental Table 1. Primer pairs and experimental conditions for RT-PCR

Supplemental Table 1. Primer pairs and experimental conditions for RT-PCR

| Gene product | Forward and reverse primers (5'–3') | Expected product size (bp) |
|---|---|---|
| AFP | TGCAGCCAAAGTGAAGAGGGAAGA<br>CATAGCGAGCAGCCCAAAGAAGAA | 217 |
| ALB | TGCTTGAATGTGCTGATGACAGGG<br>AAGGCAAGTCAGCAGGCATCTCATC | 162 |
| CYP1A2 | CAATCAGGTGGTGGTGTCAG<br>GCTCCTGGACTGTTTTCTG | 245 |
| CYP2B6 | TCCTTTCTGAGGTTCCGAGA<br>TCCCGAAGTCCCTCATAGTG | 416 |
| CYP3A4 | CAAGACCCCTTTGTGGAAAA<br>CGAGGCGACTTTCTTTCATC | 187 |
| OTC | TTTCCAAGGTTACCAGGTTACAA<br>CTGGGCAAGCAGTGTAAAAAT | 78 |
| CPS1 | CAAGTTTTGCAGTGGAATCG<br>GGACAGATGCCTGAGCCTAA | 115 |
| HNF4α | CATGGCCAAGATTGACAACCT<br>TTCCCATATGTTCCTGCATCAG | 113 |
| UBIQUITIN | GGAGCCGAGTGACACCATTG<br>CAGGGTACGACCATCTTCCAG | 118 |
| OATP1B1 | GAATGCCCAAGAGATGATGCTT<br>AACCCAGTGCAAGTGATTTCAAT | 154 |
| OATP1B3 | GTCCAGTCATTGGCTTTGCA<br>CAACCCAACGAGAGTCCTTAGG | 111 |
